# Higher resolution pooled genome-wide CRISPR knockout screening in Drosophila cells using Integration and Anti-CRISPR (IntAC)

**DOI:** 10.1101/2024.09.19.613976

**Authors:** Raghuvir Viswanatha, Samuel Entwisle, Claire Hu, Kelly Reap, Matthew Butnaru, Stephanie E. Mohr, Norbert Perrimon

## Abstract

CRISPR screens enable systematic, scalable genotype-to-phenotype mapping. We previously developed a pooled CRISPR screening method for *Drosophila melanogaster* and mosquito cell lines using plasmid transfection and site-specific integration to introduce single guide (sgRNA) libraries, followed by PCR and sequencing of integrated sgRNAs. While effective, the method relies on early constitutive Cas9 activity that potentially can lead to discrepancies between genome edits and sgRNAs detected by PCR, reducing screen accuracy. To address this issue, we introduce a new method to co-transfect a plasmid expressing the anti-CRISPR protein AcrIIa4 to suppress Cas9 activity during early sgRNA expression, which we term “IntAC” (integrase with anti-CRISPR). IntAC allowed us to construct a new CRISPR screening approach driven by the high strength *dU6:3* promoter. This new library dramatically improved precision-recall of fitness genes across the genome, retrieving 90-95% of essential gene groups within 5% error, allowing us to generate the most comprehensive list of cell fitness genes yet assembled for *Drosophila*. Our analysis determined that elevated sgRNA levels, made permissible by the IntAC approach, drove much of the improvement. The *Drosophila* fitness genes show strong correlation with human fitness genes and underscore the effects of paralogs on gene essentiality. We further demonstrate that IntAC combined with a targeted sgRNA sub-library enabled precise positive selection of a transporter under solute overload. IntAC represents a straightforward enhancement to existing *Drosophila* CRISPR screening methods, dramatically increasing accuracy, and might also be broadly applicable to virus-free CRISPR screens in other cell types, including mosquito, lepidopteran, tick, and mammalian cells.

## Introduction

CRISPR-Cas9 screens have become a critical tool for systematically mapping genotype-to-phenotype relationships [1]. In mammalian systems, these screens have been used to uncover genes essential for cell viability, immune modulation, and responses to therapeutic agents, leading to key insights into cancer biology and genetic diseases. However, adapting these powerful screens to insect models like *Drosophila melanogaster* and *Anopheles gambiae* has presented significant challenges, particularly due to the lack of an efficient retroviral induction system in these organisms. We previously showed that site-specific recombination is an efficient alternative to retroviral vector delivery for CRISPR screens and performed the first pooled CRISPR screens in insect cells [2, 3]. The approach has provided insights into essential genes and signaling pathways, hormone transport, and toxin tropism and trafficking [2, 4, 5]. Nevertheless, quantification of the precision-recall in our screening platform showed that it underperformed as compared with optimized human cell line screens, such that critical genes regulating these functions could be obscured by error or missed altogether [2].

One of the primary drawbacks in our transfection-based CRISPR screen platforms stems from the timing and control of Cas9 activity. In our previous screens, cells stably expressing Cas9 were transfected with plasmids expressing sgRNAs [2]. This poses a problem because multiple sgRNAs can be expressed from free plasmids in a single cell prior to integration into the genome, leading to genome editing, but non-integrated sgRNA plasmids decay through cell divisions, breaking the relationship between the sgRNA causing an observed phenotype and the integrated sgRNA that is detected in the next-generation sequencing (NGS) step. Therefore, we reasoned that precise temporal control of Cas9 activity, i.e., silencing activity prior to sgRNA integration, would improve screen resolution.

Strategies used to achieve temporal Cas9 control face the limitations of leakiness or low reactivation efficiency. In *Drosophila*, we previously tried treatment-inducible expression of Cas9 using a metal-inducible metallothionine promoter; however, expression of the promoter under normal media conditions was too great to suppress Cas9 editing in the uninduced state [2]. Tighter strategies for drug-controlled expression such as tet-on are lacking in *Drosophila*. Conversely, intein-based drug-inducible Cas9, while fully repressed in the non-induction condition, show low reactivation efficiency [2, 6]. Additional drug induction systems for Cas9 using TMP or rapamycin [7–9] might work in *Drosophila* but have not yet been reported. Optogenetic systems that allow for light-controlled Cas9 may offer precise control but are difficult to implement in large-scale cell culture screens [10, 11]. Introducing Cas9 in a second transfection step after sgRNA integration, the favored method in virus-free screens to date [12–14], would improve timing but introduces new challenges, such as the expense, potential for genetic drift incurred by a second large-scale transfection, and the difficulty in selecting Cas9-transfected cells for sequencing while avoiding cells that received sgRNAs but not Cas9 (which would elevate noise in the NGS step). Finally, ‘Guide-swap’, where Cas9 is electroporated into cell pools expressing sgRNAs with a dummy guide that is then ‘swapped’ for the cell-encoded guide, allows precise timing control but is expensive and challenging to scale for large screens [15].

To address these issues, we developed IntAC (integrase with anti-CRISPR), a system that integrates sgRNA libraries with built-in temporal control over Cas9 activity in a single transfection step. By expressing the anti-CRISPR protein AcrIIa4 [16–18] via a plasmid at the time of sgRNA transfection, we delay Cas9-mediated genome editing until after stable sgRNA integration has occurred. This method circumvents the need for sub-optimal induction systems or additional transfection steps, providing a straightforward and scalable solution. Another important improvement is the use of the strong *dU6:3* promoter [19]. Previous *Drosophila* CRISPR screens used the weaker *dU6:2* promoter since lowering sgRNA levels helped maintain the phenotype-sgRNA linkage [2]. The use of anti-CRISPR allowed us to use the strong *dU6:3* promoter to drive sgRNA expression in the pooled library cells, leading to higher screen resolution and improved detection of high-confidence fitness genes.

## Results

### Transient inactivation of Cas9 by anti-CRISPR

In our efforts to enhance screen resolution, we began by addressing the possibility that multiple sgRNAs are expressed in the same cell. In our previous approach, hereafter referred to as “version 1” (v.1), we transfected Cas9-expressing cells with a plasmid library of attB-flanked sgRNAs driven by the weak *dU6:2* promoter into cells that have an attP integration site (Figure 1A). The success of our previous CRISPR screens using this approach suggests that the sgRNA that integrated into the attP site predominantly led to the edit recorded at the end of the assay [2–4, 20], but it is likely that additional edits were made during the initial period following transfection. Consistent with this, pilot studies in which we increased the sgRNA levels in a small library using the stronger *dU6:3* promoter [21] led to instability of the library and death of a large fraction of transfected cells, possibly due to many cells acquiring early edits in essential genes and/or hyperactivation of the DNA damage response due to individual cells receiving too many double-strand breaks (not shown). To limit Cas9 activity until sgRNA cassette integration and loss of non-integrated sgRNA plasmids, we devised the integrase with anti-CRISPR (IntAC) approach. During library transfection, we included a plasmid encoding phage *ϕC31* integrase linked to the anti-CRISPR *AcrIIa4* by a T2A self-cleaving peptide. AcrIIa4 is the most potent known protein inhibitor of Cas9 activity [16–18]. This protein mimics guide RNAs, binding and functionally inactivating Cas9. Cas9 activity is thereby suppressed during the initial transfection period and then automatically restored when plasmids decay through a natural process of dilution during cell divisions. By this point, one sgRNA at random would have integrated into each attP site, and thus genomic edits made after anti-CRISPR activity is lost should be restricted to integrated sgRNAs (Figure 1B). Furthermore, because the IntAC approach requires only one transient transfection step and uses the highly active constitutive SpCas9, the approach improves the accuracy of genotype-phenotype mapping.

**Figure 1:**
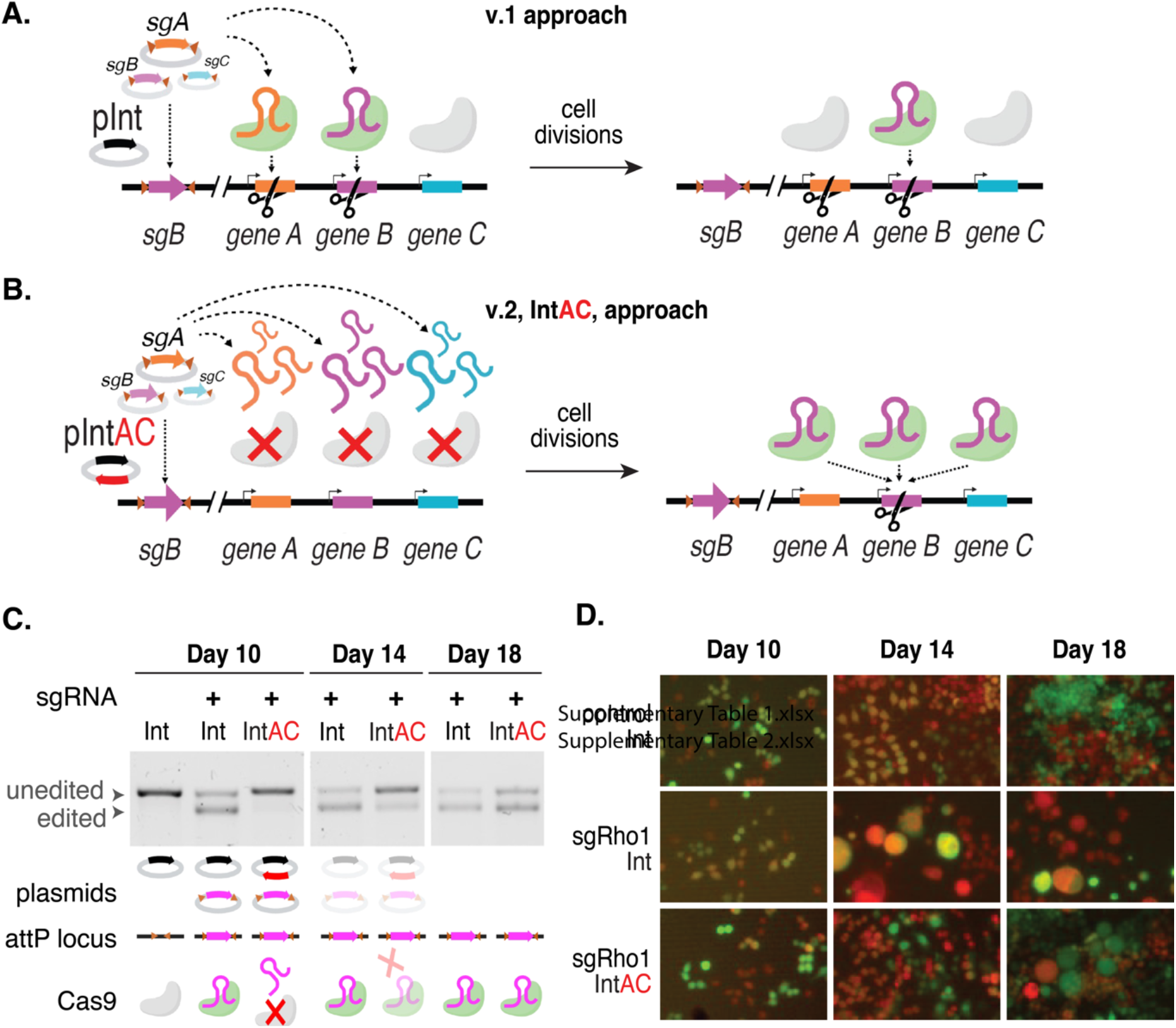
Schematic of the IntAC system and validation of transient inhibition by anti-CRISPR in *Drosophila* cells. (A,B) Schematic representation of the current CRISPR screening system compared with the IntAC system. (C) T7 endonuclease I-sensitivity assay, showing the edited versus unedited allele targeted by a CRISPR sgRNA measured over time in cells transfected as indicated. Cell populations transfected with IntAC exhibited a clear early delay in editing, with editing efficiency returning to that of controls by day 18. (D) Phenotypic validation of IntAC-delayed editing. Cells with *Rho1* suppression, characterized by isotropically overgrown cell morphology, were observed later in the IntAC population compared to controls, confirming the temporal delay in Cas9 activity.

To test whether the IntAC approach suppresses cutting by Cas9, we drove a highly efficient sgRNA targeting the *Rho1* gene with the *dU6:3* promoter and monitored editing over time using the T7 endonuclease I-sensitivity assay. Cells transfected with *pLib6.6/Rho1* alone or alongside a second non-integrating vector expressing *ϕC31* integrase (‘Int’) or *ϕC31-integrase-T2A-AcrIIa4* (‘IntAC’) were sampled at 10, 14, or 18 days after transfection. Editing reached a maximum at 10-14 days after transfection when the sgRNA vector was transfected along with Int. However, the presence of the IntAC plasmid imparted a clear delay; for the IntAC population, editing reached a maximum on day 18 (Figure 1C). As expected, the visible phenotype of *Rho1* knockout, i.e., isotropically overgrown cells [2, 22], also exhibited a delay (Figure 1D). This indicates that anti-CRISPR co-transfection imparts a delay to editing that is reversible in most cells when the plasmid encoding anti-CRISPR decays away.

### Version 2 (v.2) CRISPR screening library: optimization of sgRNA library using machine-learning and use of the strong *dU6:3* promoter

In addition to the level of editing reagents delivered to cells, previous studies have shown that optimizing sgRNA parameters through machine-learning analysis of a previous screen can also improve resolution [23]. We applied this approach, using a machine learning analysis of our previous (v.1) library screen data to develop a new ruleset for sgRNA design based on an analysis of dinucleotide position and other physical parameters, as previously described and implemented in our FlyRNAi suite of online tools [24, 25] (Figure 2A). We then constructed a new (v.2) library of 92,795 sgRNAs targeting all *Drosophila melanogaster* genes as annotated by FlyBase version 6, with a modal number of 6 sgRNAs per gene, and expressed the sgRNAs under the control of the strong *dU6:3* promoter. The library was transfected along with IntAC in two biological replicates, passaged continuously for approximately two months, and then sequenced (Figure 2B). To evaluate the library and IntAC approach, we first compared the fate of 17,948 sgRNA sequences present in both the v.1 and v.2 libraries (Figure 2C, Supplementary Table 1). In an analysis of this sgRNA set at the sgRNA level, we found that significantly more sgRNAs consistently dropped out of the pool in both replicates of v.2 as compared with v.1, suggesting that higher sgRNA levels alone led to more mutations in the v.2 cell pool as compared with v.1 (Figure 2D). To determine whether this higher level of dropout at the sgRNA level is reflected in genes detected as essential for cell fitness, the sgRNA log2 fold-changes were processed using maximum likelihood estimation (MAGeCK, [26]) to obtain gene-level Z-scores. Negative Z-scores indicate a stronger likelihood of essentiality for fitness. Using the rate of discovery of non-expressed genes (i.e., genes with an FPKM less than 1 [27]), which cannot have a role in cell fitness, to define a false discovery rate (Viswanatha et. al), we found that use of anti-CRISPR and the *dU63* promoter more than doubled the number of bona fide fitness genes that could be assigned with an error rate of 5% (Figure 2E, F, Supplementary Table 1). These results suggest that much of the improvement between v.1 and v.2 is driven by higher sgRNA expression levels via the *dU6:3* promoter as made possible by use of the IntAC approach, and that for our *Drosophila* cell system, sgRNA design optimization using machine learning had a minor impact.

**Figure 2:**
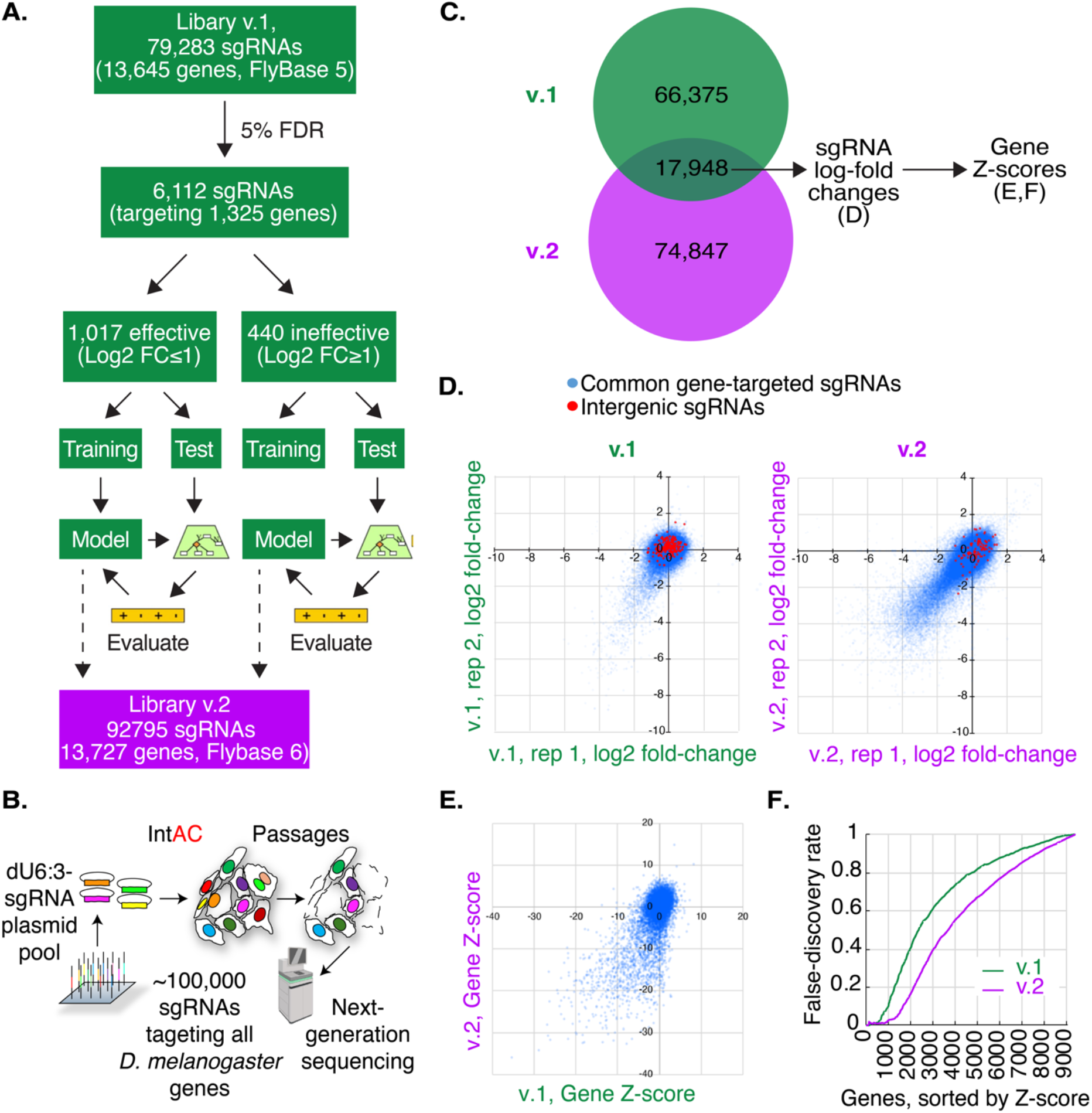
IntAC and *dU6:3* drive much of the performance increase in an updated *Drosophila* CRISPR screening library. (A) Overview of machine-learning optimization of a new (‘v.2’) genome-wide CRISPR screening library for Drosophila. (B) Schematic representation of transfection of the v.2 library. 92,795 attB-flanked sgRNAs targeting *Drosophila melanogaster* genes were transfected together. sgRNA expression was driven by the strong *dU6:3* promoter, and the library was co-transfected with the IntAC system (a plasmid expressing ϕC31 integrase and AcrIIa4) into cells. Cells were passaged for approximately two months and then CRISPR sgRNA distributions were assessed by next-generation sequencing. (C) sgRNAs common to v.1 and v.2 libraries were analyzed to isolate the effect of IntAC and *dU6:3*. (D) Comparison of 17,948 sgRNAs shows dramatically increased consistent dropout of sgRNAs in v.2 relative to v.1. (E) Gene-level analysis of the sgRNA log2-fold changes in (D) shows that the increased sgRNA dropouts translate to increased fitness gene calls (genes with negative Z-scores). (F) Cumulative distribution of false-discovery of fitness genes (genes with low expression, FPMK <1) from the gene Z-scores in (E) shows that IntAC and *dU6:3* increase the detection of fitness genes.

### IntAC screens exhibit dramatically improved fitness gene assignment

We conducted genome-wide fitness screens using the v.2 screening library with or without IntAC, and compared and the data to the v.1 screen data [2]. Compared to v.1, the v.2 screen exhibited a significantly higher number of gene dropouts, indicating improved screening efficiency. Moreover, this increase was not observed when the anti-CRISPR protein was left out (v.2 library with ϕC31 integrase alone, ‘Int’) (Figure 3A). This finding supports our hypothesis and pilot experiments suggesting that early, uncontrolled Cas9 activity in the absence of anti-CRISPR reduces the precision of the screen, particularly when combined with elevated sgRNA level due to our use of the stronger *dU6:3* promoter.

**Figure 3:**
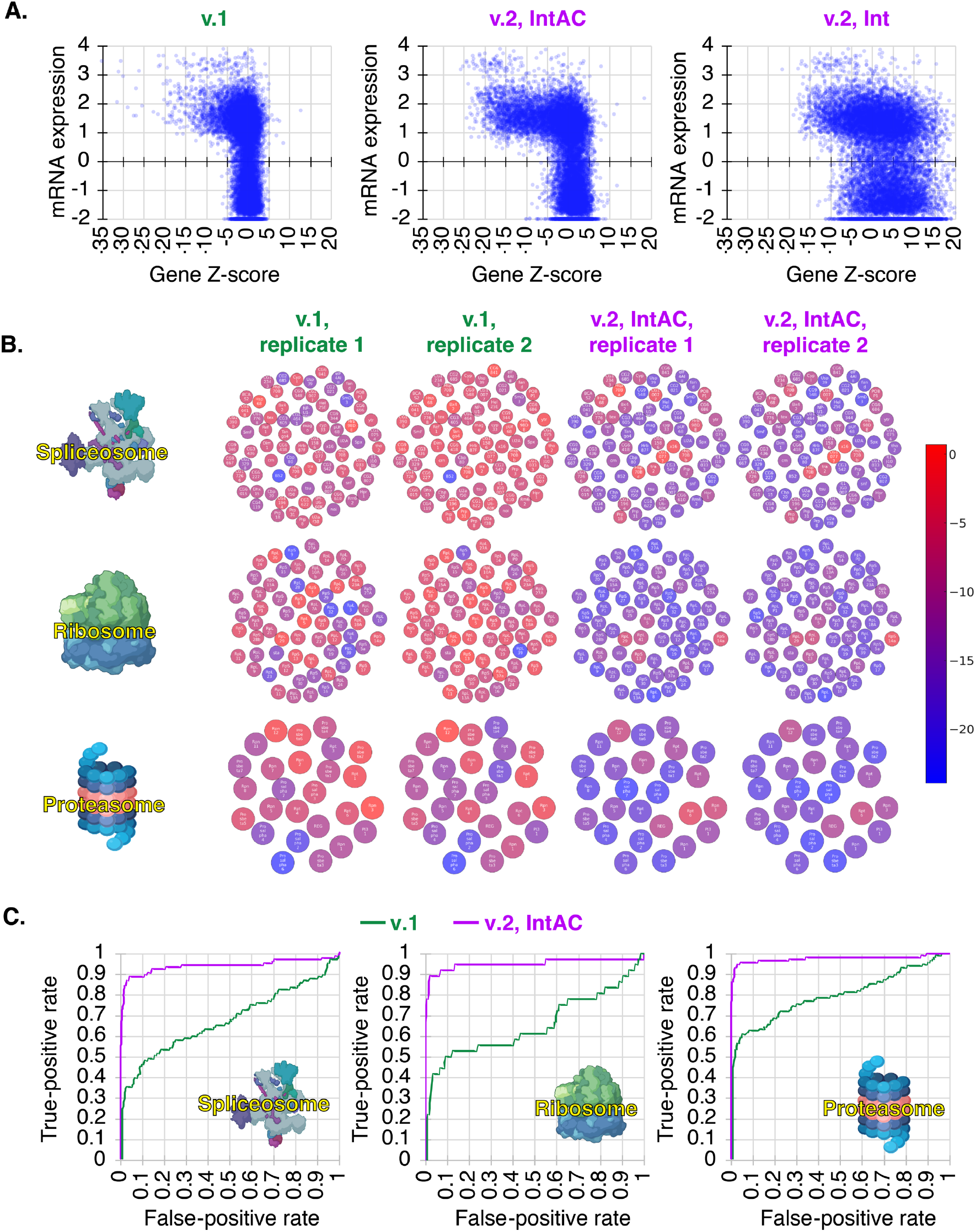
Nearly complete genome-wide cell fitness gene assignment by IntAC. (A) Comparison of gene Z-scores to mRNA expression shows that v.1 and v.2 screens identify fitness genes primarily among expressed genes, whereas Z-scores for expressed genes are lower in v.2 than v.1. Leaving out anti-CRISPR disrupts correlation between mRNA expression and Z-score, suggesting that early edits with the higher strength *dU6:3* promoter leads to instability of the library. mRNA expression is Log(FPKM+0.01). (B) Detection of essential genes across components of essential cell structures spliceosome, ribosome, and proteasome, in v.1 and v.2 screens. While these genes were enriched in v.1 screens, a substantial portion was missed. In contrast, the majority of the core components were detected in the v.2 IntAC screens, illustrating the improved sensitivity of the IntAC system for detecting essential cellular processes. The color of each component reflects its gene Z-score. (C) Precision-recall curves for fitness gene assignment in v.1 and v.2 screens, using components in (B) as true-positives and non-expressed genes as false-positives. v.2 IntAC screens exhibited a recall rate of 90-95% for essential genes, compared to 35-60% in v.1. This marked improvement in recall demonstrates that the IntAC system yields more comprehensive and accurate identification of fitness genes across the genome.

To determine whether known essential genes were detected as fitness genes, we asked whether core components of the spliceosome, ribosome, or proteasome (as assigned by KEGG) were detected in gene-level analysis of v.1 or v.2, IntAC, screens (Supplementary Table 1). While these gene sets were statistically enriched in v.1 screens, the majority of their component genes were missed. Conversely, in v.2 IntAC screens, most genes in each gene set were detected (Figure 3B). Moreover, as expected, comparison of the two gene sets shows that the v.1 fitness gene set is a subset of the v.2 fitness gene set (Supplementary Figure 1). To quantify the rate of recall assuming that the KEGG assignment is correct, we assigned the genes in each KEGG gene set as the set of true positives. Using non-expressed genes (FPKM <1) as the set of false positives, we estimated a precision-recall rate. The results show that the recall at 5% is 35-60% for v.1 and 90-95% with v.2 (Figure 3C). Thus, using the improved v.2 IntAC CRISPR screen approach, we identified the most complete and accurate set of cell fitness genes yet assembled for *Drosophila* or any invertebrate, likely capturing 90-95% of all fitness genes with 5% error. Importantly, we note that this approach and library has now been applied independently by multiple members of our group and resulted in comparable precision-recall rates (Suppl. Fig. 2A,B).

### An updated comparison of *Drosophila* and human cell fitness genes

We next compared the high-resolution *Drosophila* cell fitness screen data to data from human cell line CRISPR screens. First, we determined the *Drosophila* orthologs of human ‘core essential genes’ (CEG2) assembled from human CRISPR and RNAi screens [28]. The majority of these genes are present in our *Drosophila* v.2 CRISPR screen (∼90% overlap allowing 5% false-discovery in the *Drosophila* v.2 CRISPR screen data, Supplementary Figure 3), but CEG2 orthologs only accounts for ∼17% of the *Drosophila* v.2 geneset. We reasoned that some of the remaining fitness genes in *Drosophila* cells might reflect genes that were missed due to masking of essential gene functions in human CRISPR screens. Whole-genome duplication (WGD) events create paralogs with the potential for redundant functions; invertebrates underwent fewer WGD events than humans [29]. To address this, we asked whether our CRISPR screen data correlates with human cell line CRISPR screens for *Drosophila* genes with a 1-to-1 relationship with human genes, and whether our findings for “1-to-many” genes lends further support to the idea that increased genetic redundancy in human cells masks detection of essential gene functions. To do this, we first retrieved all human gene orthologs for all *Drosophila* genes using the DIOPT approach [30] and then computed the first quartile of all Cas9-mediated Essentiality Ranking Evaluator of Specificity (CERES) scores (‘Q1 CERES’) to represent a consensus fitness score across all human cell lines for every gene tested in the DepMap [31]. To map orthologs, we used a DIOPT score of 6 or greater, resulting in 4,380 1-to-1 orthologs and 2,206 1-to-many orthologs. We observed a strong correlation for 1-to-1 orthologs in both species (R^2^ = 0.41, Figure 4A). However, consistent with expectation, this correlation fell significantly for genes that have two or more orthologs in humans (R^2^ = 0.18), even when we intentionally counter-biased the analysis by choosing the most essential human ortholog to represent the human gene in the pair (Figure 4B). This finding supports the idea that redundancy due to paralogs is an important factor in CRISPR screen data interpretation, and highlights paralog groups in human cells that do not score positive in screens but collectively represent an essential function, as their unique Drosophila ortholog is required for optimal fitness. A list of 123 gene groups matching these criteria is provided in Supplementary Table 2.

**Figure 4:**
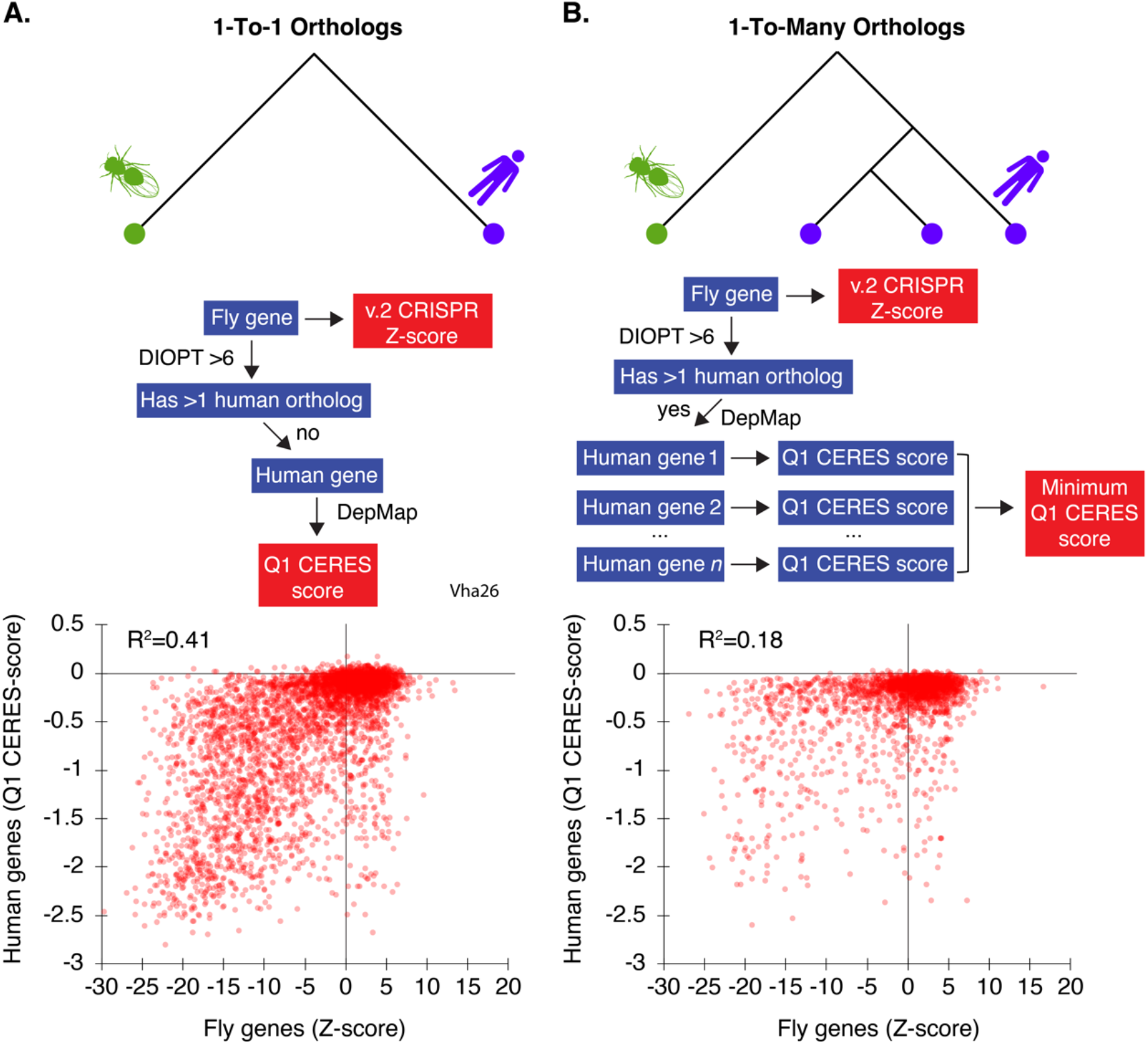
Correlation between *Drosophila* fitness genes and human orthologs in IntAC screens. (A) For genes with a 1-to-1 ortholog relationship between *Drosophila* and humans, a strong correlation (R² = 0.41) was observed, suggesting high conservation of fitness genes across species. Schematic of processing steps comparing *Drosophila* orthologs to their predicted human orthologs (assessed by a DIOPT score >6, using the DRSC Integrative Ortholog Prediction Tool [30]). Genes with a 1-to-1 relationship were retained for analysis, and the first-quartile (Q1) of their Cas9-mediated Essentiality Ranking Evaluator of Specificity (CERES) scores was plotted across all DepMap data (23Q4) [31]. (B) Correlation analysis for *Drosophila* genes with multiple orthologs in humans. In cases where *Drosophila* genes had multiple human orthologs (1-to-many relationships), the correlation with fitness scores dropped significantly (R² = 0.18). Schematic showing processing steps. When multiple human orthologs had a DIOPT score > 6, the ortholog with the lowest CERES score was chosen in order to counter-bias the data against selecting an inactive paralog. This result suggests that paralog redundancy in humans can mask the essentiality of certain genes.

### Positive selection of transporters mediating solute overload using IntAC screens

Next, we asked how the IntAC method performs in positive-selection screens and with small focused sub-libraries of sgRNAs. Matching solutes to their transporters is important to understand how cells manage intake, export, and compartmentalization of solutes and how this process goes awry in metabolic disorders [32]. In *Drosophila*, approximately one third of predicted transporters are annotated as “CGs” (computed genes) without documented phenotypes [30]. To provide greater insight into these genes, we set out to target them with higher resolution than the genome-wide library i.e., using more sgRNAs per gene. We prepared a new sub-library containing 10 sgRNAs per gene for 389 solute carrier genes, selecting only those expressed in *Drosophila* S2R+ cells. We then transfected cells using the IntAC approach (Figure 5A). In a pilot panel of solutes, we discovered that the nucleoside cytidine was extremely toxic to cells at a concentration of 250 µM (not shown). We then performed two biological replicates of a screen for resistance to cytidine overload. The genes *Ent1* and *Ent2* are thought to mediate nucleoside uptake in *Drosophila* [33, 34], but only *Ent2* is expressed in S2R+ cells [35]. Following selection, *Ent2* sgRNAs predominated reads, and non-*Ent2* sgRNAs (likely passengers) emerged only after the seventh *Ent2* sgRNA in replicate 1 or the fifth *Ent2* sgRNA in replicate 2 in the ranked dataset (Figure 5B, C; Supplementary Table 1). The precision of this selection again suggests that IntAC results in cells edited primarily by integrated sgRNAs, even when in this case we combine it not only with the use of higher strength promoter *dU6:3*, but also use of a positive selection assay and a smaller sgRNA library. Thus, the IntAC approach is extensible to positive selection screens and focused sub-libraries.

**Figure 5:**
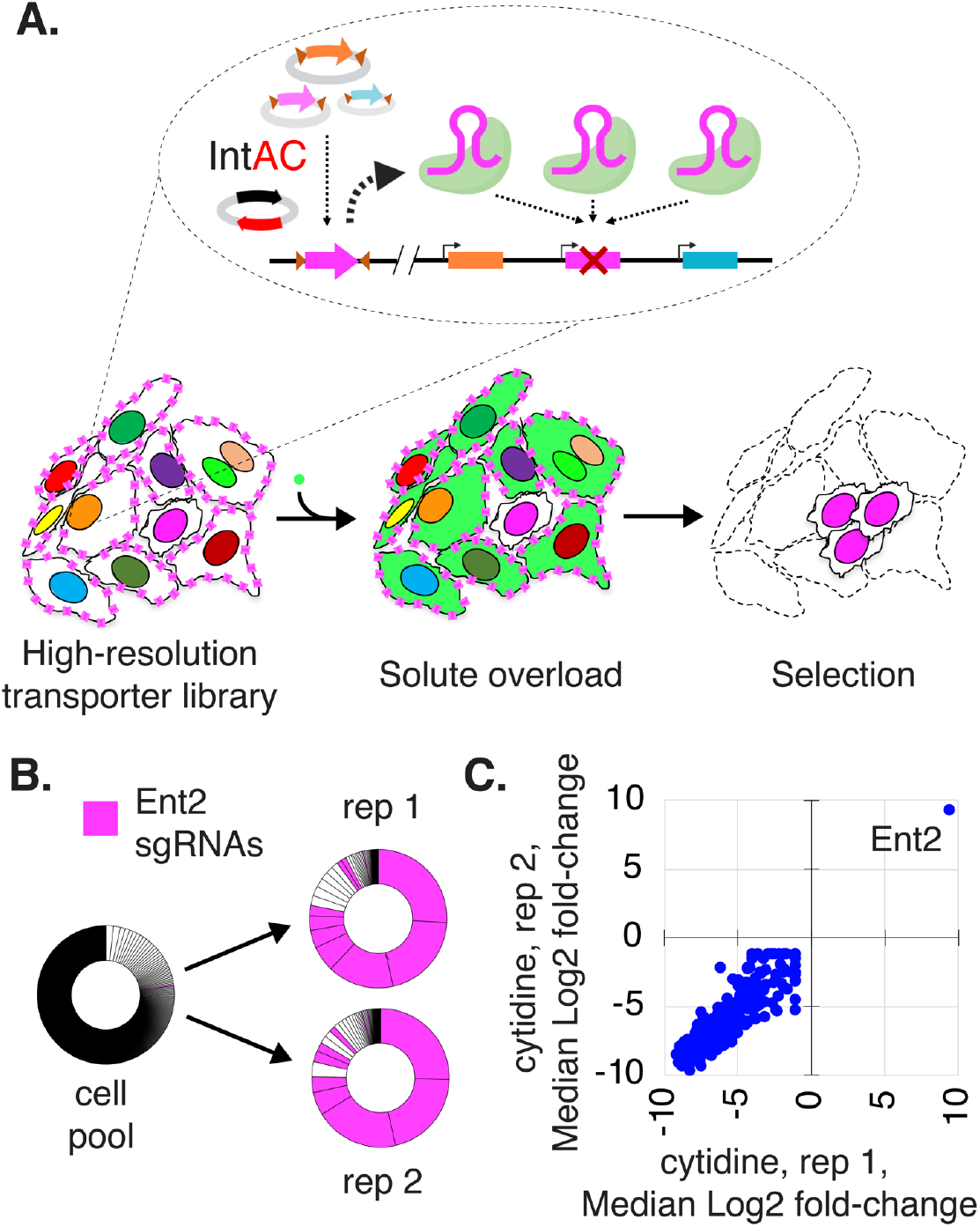
Positive selection of a nucleoside transporter using IntAC in a solute overload screen. (A) Schematic of a solute overload screen using the IntAC platform. Cells were transfected with a targeted sublibrary of sgRNAs focused on 389 predicted solute carrier genes, using the *dU6:3* promoter to drive sgRNA expression. Following transfection, the cells were subjected to solute overload, specifically with adenosine and cytidine, to identify genes involved in nucleoside transport and cellular resistance to solute toxicity. (B) Representation of sgRNA distribution following cytosine overload. All ∼4,000 sgRNAs in the library are represented as slice of the donut chart in descending abundance from 12:00 clockwise. Following two independent overloads with cytosine, different *Ent2* sgRNA sequences (pink) occupy ∼75% of reads and the first 7 positions of replicate 1 and the first 5 positions of replicate 2. (C) Gene-level Z-scores show that *Ent2* is the only gene detected in both replicates.

## Discussion

CRISPR pooled screening is the state-of-the-art for genetic screening in *Drosophila* cells [2]. The method has already provided insights into essential genes and signaling pathways, hormone transport, pathogen tropism, and toxin trafficking [2, 20, 36]. Nevertheless, the measured efficiency left room for improvement, especially when compared to the precision-recall achieved by optimized mammalian cell CRISPR screens [23]. Here, we explored whether temporal control over Cas9 activity could achieve better resolution. We introduced a straightforward method to suppress unwanted Cas9 activity prior to integration of sgRNAs during recombination-based CRISPR screens, the IntAC approach.

In our efforts to enhance screen resolution, we introduced two key modifications at the same time: the use of the stronger *dU6:3* promoter to drive sgRNA expression and optimization of sgRNA designs using a machine learning approach based on our previous screens. By examining sgRNAs common to both the original and machine learning-based sgRNA design sets, we were able to observe a significant performance improvement due to IntAC and *dU63*, but we were unable to quantify the contribution of guide optimization *per se*. Formally, this would require construction of a new CRISPR library driven by *dU6:3* using the full set of v.1 sgRNAs. Due to the significant time required for new library construction and minimal information we anticipated gaining by testing such a library, we deemed this outside of the scope our current work. Based on work from others [23], we do anticipate that application of a machine learning approach can improve library design. In the future, we plan to use machine learning to evaluate sgRNA design rules based on the new, optimized v.2 screen platform data, and further optimize screen parameters based on the outcome. We also plan to test the impact of modifications to guide architecture that have proven beneficial for mammalian CRISPR screens [37].

Using IntAC along with a new *dU6:*3-driven sgRNA library enhanced resolution, resulting in the most comprehensive list of cell essential genes in *Drosophila*. Although some of these genes are likely specific to the *Drosophila* cell type studied, i.e., S2R+ cells, which are thought to have originated from hemocyte precursors [38, 39], many of the genes identified in the screen are likely to reflect factors necessary for growth and survival of all cells. Consistent with this, 90% of the previously defined ‘core essential geneset 2’ (CEG2) derived from mammalian screen data can be found within our *Drosophila* screen. Interestingly, in addition, comparison of the *Drosophila* cell screen data to equivalent data from human cells allowed us to predict human genes whose evolutionary duplication might mask their importance. Studies in *Drosophila* cells could help clarify the role of these crucial gene families in both invertebrates and mammals. This work underscores the utility of straightforward single-gene knockout screens in *Drosophila* cells to identify genes that would be missed in similar screens in mammalian cells.

We also show that IntAC displays precision in positive selection screens. In these experiments, we developed the first focused CRISPR library targeting *Drosophila* solute transporters, enabling the identification of transporters responsible for solute overload using smaller cell populations than are needed to conduct gnome-wide screens. This will enable the creation of comprehensive solute-to-transporter maps and provides a platform for *in vivo* follow-up studies in *Drosophila*. Such insights can be crucial for understanding transport mechanisms, including homologous processes in human cells.

Given the robust performance of the IntAC approach in *Drosophila* cells, a next goal is to apply the approach in other cell species. This could have particular impact on non-model species for which cell lines exist, but the application of large-scale genetic screening would be novel. This includes cell lines derived from lepidopteran insect pests and arthropod vectors of disease such as mosquitos and ticks. Given the lack of any functional genomic data, genome-wide screens enhanced by IntAC could be used to construct improved gene annotations and reveal important evolutionary insights of cell essential genes. Furthermore, the identification of essential genes in mosquitos and ticks can be used as targets for gene drives designed for population suppression [40, 41]. The IntAC method relies on the loss of the anti-CRISPR-encoding plasmids over time. At low frequency, however, plasmids can integrate into the cell genome, independent of site-specific integration of sgRNA cassettes. The IntAC method relies on the loss of the anti-CRISPR-encoding plasmids over time. At low frequency, however, these plasmids can integrate into the cell genome, independently of the site-specific integration of sgRNA cassettes. If the efficiency of site-specific integration is low, then the likelihood of selecting drug-resistant cells with plasmid sequences randomly integrated is relatively high. This includes anti-CRISPR plasmids which would permanently inactivate Cas9 (such that editing would not occur). The sgRNAs in such cells would nevertheless remain part of the cell pool, weakening the genotype-phenotype linkage. Thus, the balance between site-specific and random integration in any new cell systems will have to be tested and optimized. Note that in the PT5 S2R+ cell line we used in this study, we estimated that fewer than 1% of drug-resistant cells have randomly integrated plasmids [2]. Second, the method relies on the availability of efficient anti-CRISPRs for the Cas enzyme utilized. Thus, IntAC is amenable in principle to all screens using Cas9 or Cas12a (inhibited by the AcrVA family [42]) including kinase-dead or nicking variants for CRISPR activation or inhibition [1], prime editing [43], or base editing [44], whereas there is currently no anti-CRISPR for the RNA-cleaving Cas13d/CasRx. Thus, while the method is widely applicable to many currently used screening types, the universality will depend on the development of precise anti-CRISPRs for each Cas. Finally, IntAC could also potentially enhance mammalian virus-free CRISPR screens which have similarly used a plasmid transfection-based attP-attB recombination to deliver sgRNAs followed by a second transfection to deliver Cas9. These have been demonstrated in CHO, HEK293, and K562 cells [13, 45–48], where the key motivations were to design CRISPR screens that expressed constant amounts of sgRNAs from a defined locus and to avoid biosafety constraints of using viral vectors. In such cases, IntAC could offer a promising addition by enabling temporal control of Cas9 activity in a single transfection step and lead to improved screening resolution.

## Materials and Methods

### Cell lines and treatments

The *Drosophila* S2R+ derivative PT5 (NPT005; DGRC #229) was transfected with pMK33/Cas9 [2] or with pDmAct5C::Cas9-2A-Neo [25] and maintained in either 200 ng/ µL Hygromycin B (Calbiochem) or 500 µg/mL G-418 (Goldbio). The efficiency of CRISPR sgRNAs in each cell line was identical (not shown), and the cell lines were used interchangeably for these experiments. Cytidine (Sigma, C122106) was dissolved in Schneider’s media to prepare a 2x stock to achieve a final concentration of 250 µM. Images were captured using an Evos 5000 Microscope equipped with a 20X objective and GFP and Texas Red filters.

### Plasmids

pLib6.6 (Addgene # 176652) was constructed by replacing the *dU6:2* promoter from pLib6.4 (Addgene #133783) with the *dU6:3* promoter from pCFD3 (Addgene #49410, [19]). A synthetic gene fragment containing the anti-CRISPR AcrIIa4 codon-optimized for *Drosophila* was attached downstream of the ϕC31[2] integrase (from pBS130, Addgene # 26290) followed by a P2A site. The resulting fusion was cloned downstream of the *Drosophila* Actin promoter in pAWF using standard cloning methods to generate pIntAC (which will be made available on Addgene shortly). As a control, pInt was generated for this study by cloning ϕC31 integrase (from pBS130, Addgene # 26290) into pAWF.

### T7 Endonuclease-I assay

A pLib6.6 plasmid expressing an sgRNA targeting Rho1 was used alongside a plasmid encoding ϕC31 integrase (Int) or ϕC31 integrase-2A-ActIIa4 (IntAC) under the *Drosophila* Actin promoter. The *Rho1* target genomic region was amplified by PCR using specific primers flanking the CRISPR/Cas9 editing site (m-Rho1-seq20_0F, 5’-GGT GCC TGC GGT AAA ACT TG-3’ and dm-Rho1-seq20_0R, 5’- ATC ACT TGG ATG GCA GGG TG-3’). The PCR products were gel extracted and then denatured at 95°C for 5 minutes and re-annealed by gradually cooling to room temperature to form heteroduplex DNA. The re-annealed DNA was then digested with 10 units of T7 Endonuclease I (New England Biolabs, Cat# M0302) in 1X NEBuffer 2 at 37°C for 30 minutes. The reaction was stopped by adding 0.25 M EDTA, and the products were analyzed on a 2% agarose gel stained with SYBR Safe DNA stain (Thermo).

### CRISPR library construction and delivery

CRISPR libraries were synthesized as oligo pools between constant sequences (5’ sequence: TAT ATA GAC CTA TTT TCA ATT TAA CGT CG; 3’ sequence: GTT TTA GAG CTA GAA ATA GCA AGT TAA AAT) (Genscript), amplified with outside primers corresponding to the 5’ constant sequence and the reverse complement of the 3’ constant sequence in 17 cycles using Phusion Polymerase (New England Biolabs), and then inserted into BbsI-digested pLib6.6 using the NEB HiFi Assembly Master Mix Library.The reaction was electroporated into E. cloni 10G ELITE cells (Lucigen) and plated on 150-mm LB-Carbenicillin selective plates and grown overnight for 30°C. This procedure was scaled such that the number of resulting colonies was at least 100 times the library complexity. Transfection in *Drosophila* S2R+ derivative cell lines was performed as previously described [2] using Effectene reagent along with minor adjustments for cost savings (detailed in Supplementary Figure 2A). All libraries were transformed with pInt or pIntAC at a molar ratio of 1:1. Briefly, 5 µg of library and 5 µg of either pInt or pIntAC was mixed with 80 µL of Enhancer Solution, 1500 µL of EC buffer, and 300 µL of Effectene (Qiagen). Transfection mix was added to 50 mL of actively growing S2R+ cells adjusted to 2.4 x 10^6^ cells/mL, mixed, and 5 mL distributed to each of ten 100-mm dishes and shaken to distribute the cells evenly. Plates were tightly sealed in plastic film and incubated overnight at room temperature. The next day, 5 mL of fresh media was added to each plate to prevent evaporation. After three additional days, cells were selected in puromycin-containing media. They were then grown for 3 additional weeks under puromycin selection with media changes every 4-5 days before being used for subsequent experiments. For chemical selection experiments, chemicals were added at this point. For fitness gene assessment, cells were passaged for an additional month using the same regimen.

### Library sequencing and CRISPR screen data analysis

The sgRNA counts were prepared using a genomic DNA PCR protocol established previously [2] with minor modifications. First, cells were lifted from plates and resuspended at a concentration of 5-20 x 10^6^ per mL, and 25-50 mL of cells was pelleted in a 50 mL conical tube and the media discarded. Pellets were stored frozen. Next, eight Zymo gDNA miniprep (D3025) columns were used per frozen pellet using the manufacturer’s instructions, resulting in the isolation of 300-400 µL of gDNA per sample. Samples were diluted 1:1 with water and again 1:1 with 2X GoTaq Master Mix (Promega M712B) containing appropriate primers and amplified in 23 PCR cycles. Samples were resolved on a 1% agarose gel and the ∼450 bp band corresponding to sgRNA expression cassette containing a mixture of sgRNA sequences was gel purified. Next outside primers adding P5 and P7 sequences were used to reamplify the product using Phusion Polymerase (New England Biolabs). The concentration of DNA in each sample was next quantified using a Cubit fluorometery assay (dsDNA, Broadrange, Thermo) and then normalized and mixed together. Combined samples were then directly loaded on a Novaseq 6000 lane (Biopolymers Facility, Harvard Medical School), and sequencing was conducted with the goal of sequencing each sgRNA 100 times or more. All data processing used MAGeCK version 0.5.6.

### Barcoding strategy

We used a two-step fingerprinting-and-barcoding strategy similar to that previously employed [2]. A portion of the Illumina Read1 primer, unique experimental fingerprints (varying stretches of random nucleotides), and 6-nucleotide barcodes and were added to the first PCR round (PCR1). The second PCR (PCR2) then anneals to the PCR1 product and adds extensions containing the 5’ P5 site and 3’ P7 site to enable sequencing on Illumina sequencers. An example of one forward primer used for the first PCR step is: 5’- CCT ACA CGA CGC TCT TCC GAT CTN NNN TTG GCT CCT ATT TTC AAT TTA ACG TCG-3’, where “NNNN” represents the fingerprint and “TTGGCT” represents the barcode. “CCT ATT TTC AAT TTA ACG TCG” aligns to the dU6:3 primer and extends the unique sgRNA spacers. The common PCR1 reverse primer is 5’-TTT GTG TTT TTA GAA TAT AGA ATT GCA TGC-3’. The common PCR2 forward primer (adding P5) is: 5’-AAT GAT ACG GCG ACC ACC GAG ATC TAC ACT CTT TCC CTA CAC GAC GCT CTT CCG ATC T-3’. The common PCR2 reverse primer (adding P7) is: 5’-CAA GCA GAA GAC GGC ATA CGA GAT TTT GTG TTT TTA GAA TAT AGA ATT GCA TGC-3’.

## Acknowledgements

We thank Baolong Xia, Ahmad Muhammad, and Pratyajit Mohapatra for testing the method. This work was supported by NIH P41 GM132087 to SEM. and NP. NP is a Howard Hughes Medical Institute (HHMI) investigator. This article is subject to HHMI’s Open Access to Publications policy. HHMI lab heads have previously granted a non-exclusive CC BY 4.0 license to the public and a sublicensable license to HHMI in their research articles. Pursuant to those licenses, the author-accepted manuscript of this article can be made freely available under a CC BY 4.0 license immediately upon publication.

---

**Supplementary Figure 1:**
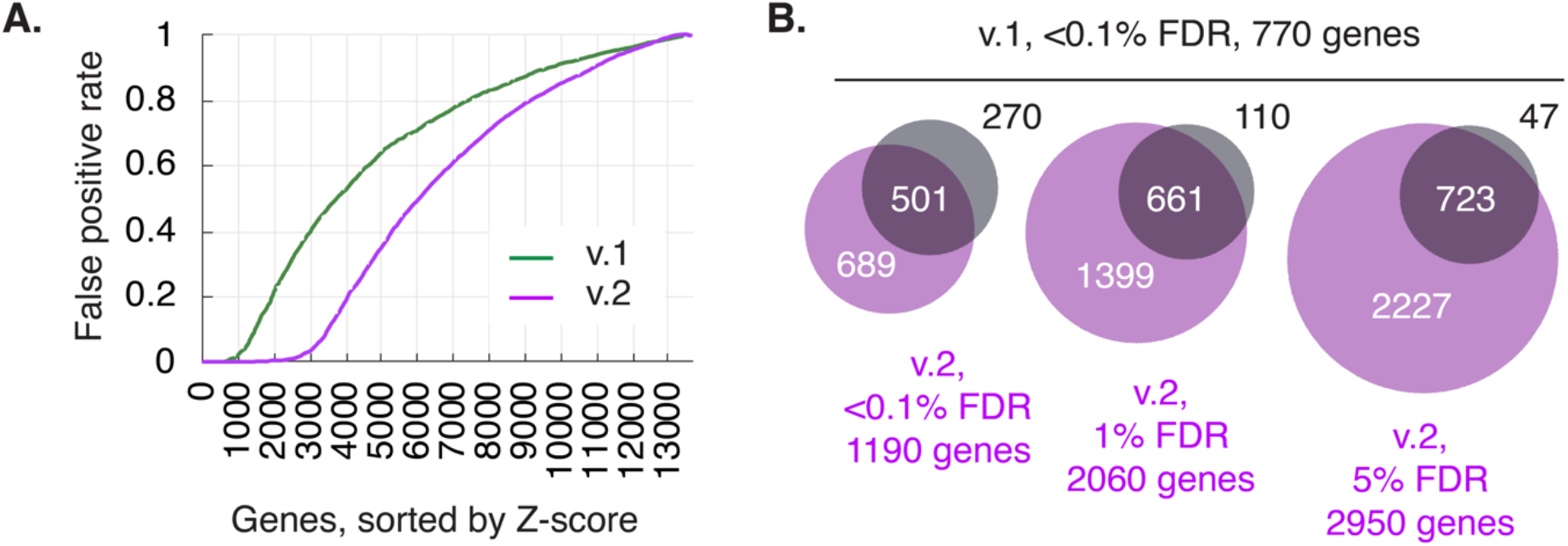
Comparison of v.1 and v.2 screens. V.2 screens produce a larger set of fitness genes compared with v.1. (A) Cumulative distribution of false-positive rate (using non-expressed genes as false-positives, FKPM<1) as a function of genes, sorted in ascending value by gene Z-score (greater-to-less likelihood of being essential for cell fitness). (B) The 770 genes with the strongest likelihood of representing bona fide fitness genes in v.1 screens are mostly a subset of those detected in v.2. The overlap becomes progressively clearer when the false-discovery rate (FDR) of v.2 fitness genes is relaxed.

**Supplementary Figure 2:**
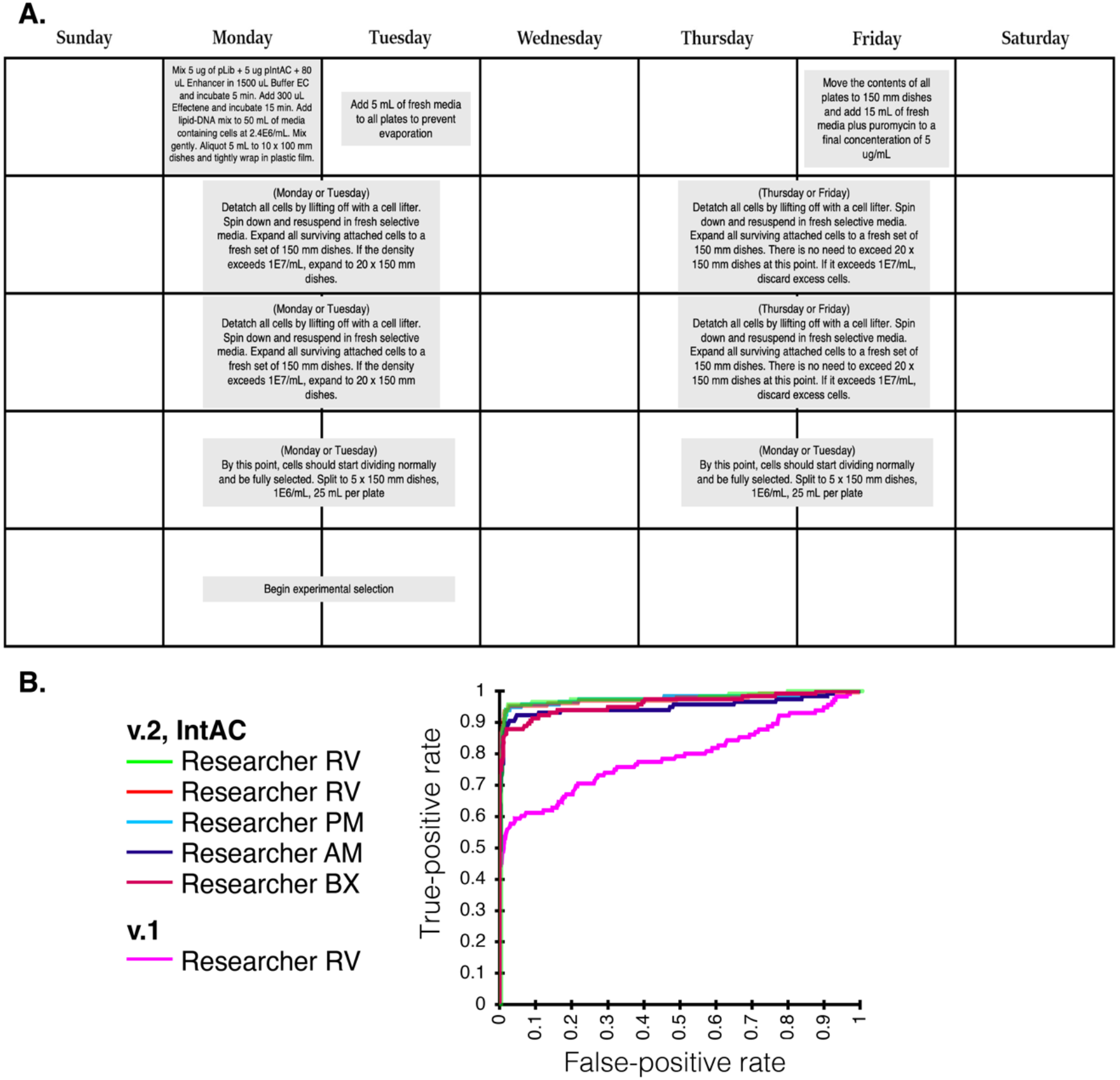
Validation of IntAC CRISPR screen across independent trials and researchers. (A) Hands-on steps for generating a pooled library using IntAC. (B) Performance of the IntAC platform in independent trials by different researchers. Data from multiple independent screens conducted using the IntAC platform shows high reproducibility in terms of sgRNA dropout patterns and gene Z-scores. A precision-recall analysis comparing the detection of essential genes (true-positives, KEGG-assigned ribosome and proteasome genes) versus false-positives (non-expressed genes, genes with mRNA FPKM < 1). These independent trials confirmed the high precision and robustness of the v.2, IntAC, system relative to the previous v.1 CRISPR screening system in the hands of different researchers

**Supplementary Figure 3:**
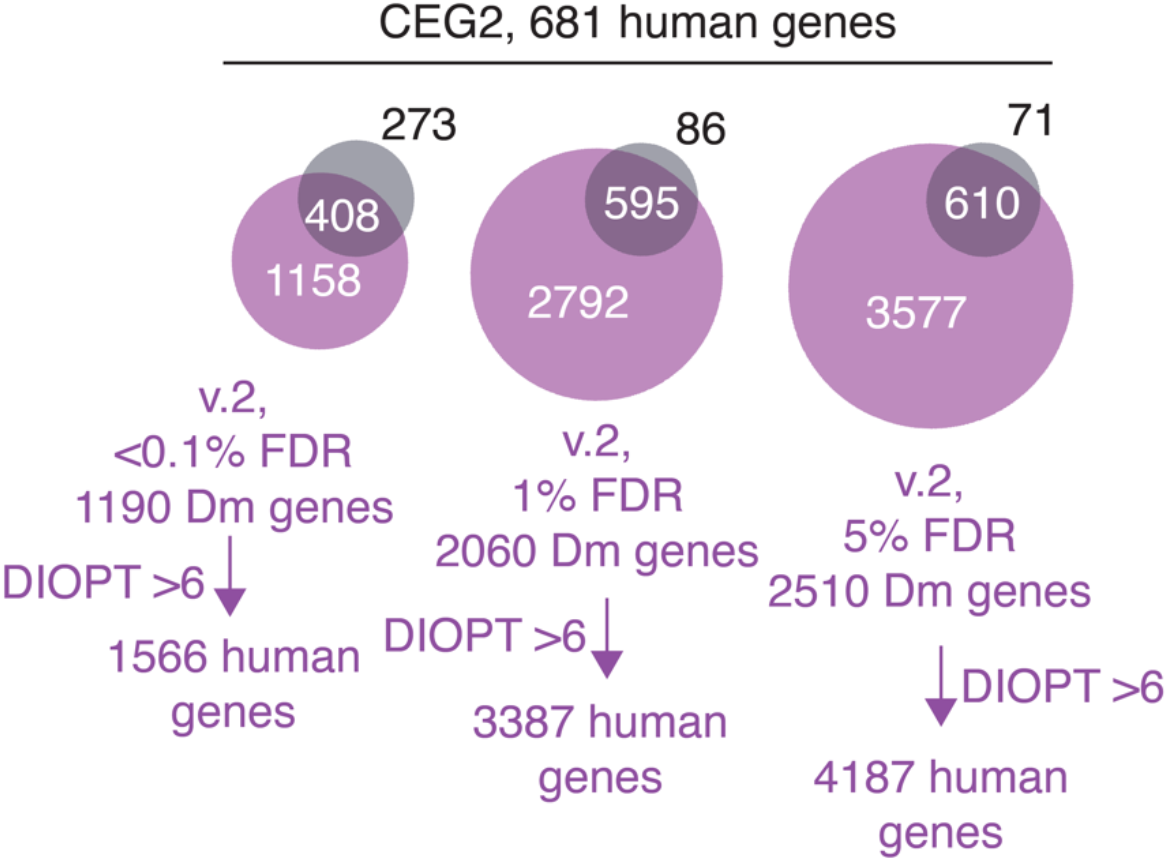
Overlap between human cell essential geneset and *Drosophila* cell essential geneset. The 681 genes in the core essential geneset v.2 [49]. The overlap becomes progressively clearer when the false-discovery rate (FDR) of v.2 fitness genes is relaxed.

**Supplementary Table 1.** Raw and processed data for IntAC CRISPR screens. The ‘sgRNA-level’ tabs contain the unprocessed count data from two different screens. For the fitness screen, the count data was analyzed using the MAGeCK ‘maximum likelihood estimation (MLE)’ package to calculate Z-scores. These Z-scores indicate the likelihood that a given gene was enriched (positive Z-score) or depleted (negative Z-score) in the cell pools [26]. The results are presented in the ‘gene-level’ tab. Additionally, mRNA expression data for each gene, as compiled by modENCODE [27], is provided. For the cytidine overload screen, we used the ‘test’ package in MAGeCK [26] to calculate the median log2 fold-change for the 10 sgRNAs targeting each gene. This information is also included in the ‘gene-level’ tab.

**Supplementary Table 2.** Putative fitness gene groups in human cells identified by the essentiality of their unique *Drosophila* ortholog. We present 123 genes scoring as essential for *Drosophila* cell fitness (false-discovery rate < 5%) for which the putative human orthologs (DIOPT score > 6 [30]) do not score as essential for human cell fitness. For this analysis, we report only gene families for which the minimum first quartile CERES value is greater than -0.5, indicating that none of the human orthologs is strongly essential for cell fitness in the DepMap [31].

## Notes

### Competing Interest Statement

The authors have declared no competing interest.

### Summary of Updates

We updated the manuscript to include minor changes to the text, additions to the authors list, and corrections to supplementary figures.

## References

1. Villiger, L., et al., CRISPR technologies for genome, epigenome and transcriptome editing. Nature Reviews Molecular Cell Biology, 2024. 25(6): p. 464–487.

2. Viswanatha, R., et al., Pooled genome-wide CRISPR screening for basal and context-specific fitness gene essentiality in Drosophila cells. eLife, 2018. 7.

3. Viswanatha, R., et al., Bioinformatic and cell-based tools for pooled CRISPR knockout screening in mosquitos. Nature Communications 2021 12:1, 2021. **12**(1): p. 1-13.

4. Okamoto, N., et al., A Membrane Transporter Is Required for Steroid Hormone Uptake in Drosophila. Developmental cell, 2018. 47(3): p. 294–305.e7.

5. Xu, Y., et al., Genome-wide CRISPR Screens in Drosophila Cells Identify Vsg as a Tc Toxin Receptor. Nature. (In review), 2022.

6. Davis, K.M., et al., Small molecule–triggered Cas9 protein with improved genome-editing specificity. Nature Chemical Biology, 2015. 11(5): p. 316–318.

7. Zetsche, B., S.E. Volz, and F. Zhang, A split-Cas9 architecture for inducible genome editing and transcription modulation. Nature Biotechnology, 2015. 33(2): p. 139–142.

8. Maji, B., et al., Multidimensional chemical control of CRISPR–Cas9. Nature Chemical Biology, 2016. 13(1): p. 9–11.

9. Richter, F., et al., Switchable Cas9. Current Opinion in Biotechnology, 2017. 48: p. 119–126.

10. Nihongaki, Y., et al., Photoactivatable CRISPR-Cas9 for optogenetic genome editing. Nature Biotechnology, 2015. 33(7): p. 755–760.

11. Bubeck, F., et al., Engineered anti-CRISPR proteins for optogenetic control of CRISPR– Cas9. Nature Methods, 2018. 15(11): p. 924–927.

12. Chang, J., et al., Genome-wide CRISPR screening reveals genes essential for cell viability and resistance to abiotic and biotic stresses in Bombyx mori. Genome Research, 2020. 30(5): p. 757–767.

13. Xiong, K., et al., An optimized genome-wide, virus-free CRISPR screen for mammalian cells. Cell Reports Methods, 2021. 1(4): p. 100062.

14. Kim, S.H., et al., Identification of hyperosmotic stress-responsive genes in Chinese hamster ovary cells via genome-wide virus-free CRISPR/Cas9 screening. Metab Eng, 2023. 80: p. 66–77.

15. Ting, P.Y., et al., Guide Swap enables genome-scale pooled CRISPR–Cas9 screening in human primary cells. Nature Methods, 2018. 15(11): p. 941–946.

16. Dong, D., et al., Structural basis of CRISPR–SpyCas9 inhibition by an anti-CRISPR protein. Nature, 2017. 546(7658): p. 436–439.

17. Shin, J., et al., Disabling Cas9 by an anti-CRISPR DNA mimic. Science Advances, 2017. 3(7): p. e1701620.

18. Yang, H. and D.J. Patel, Inhibition Mechanism of an Anti-CRISPR Suppressor AcrIIA4 Targeting SpyCas9. Molecular Cell, 2017. 67(1): p. 117–127.e5.

19. Port, F., et al., Optimized CRISPR/Cas tools for efficient germline and somatic genome engineering in Drosophila. Proceedings of the National Academy of Sciences, 2014. 111(29): p. E2967–E2976.

20. Xu, Y., et al., Genome-wide CRISPR Screens in Drosophila Cells Identify Vsg as a Tc Toxin Receptor. Nature (in review).

21. Port, F., et al., Optimized CRISPR/Cas tools for efficient germline and somatic genome engineering in Drosophila. Proceedings of the National Academy of Sciences, 2014. 111(29): p. E2967–E2976.

22. Rogers, S.L. and G.C. Rogers, Culture of Drosophila S2 cells and their use for RNAi-mediated loss-of-function studies and immunofluorescence microscopy. Nature Protocols, 2008. 3(4): p. 606–611.

23. Doench, J.G., et al., Optimized sgRNA design to maximize activity and minimize off-target effects of CRISPR-Cas9. Nature Biotechnology, 2016. 34(2): p. 184–191.

24. Hu, Y., et al., FlyRNAi.org—the database of the Drosophila RNAi screening center and transgenic RNAi project: 2021 update. Nucleic Acids Research, 2021. 49(D1): p. D908–D915.

25. Viswanatha, R., et al., Bioinformatic and cell-based tools for pooled CRISPR knockout screening in mosquitos. Nature communications, 2021. 12(1).

26. Li, W., et al., MAGeCK enables robust identification of essential genes from genome-scale CRISPR/Cas9 knockout screens. Genome biology, 2014. 15(12): p. 554–554.

27. Roy, S., et al., Identification of Functional Elements and Regulatory Circuits by Drosophila modENCODE. Science, 2010. 330(6012): p. 1787–1797.

28. Hart, T., et al., Evaluation and Design of Genome-Wide CRISPR/SpCas9 Knockout Screens. G3 (Bethesda, Md.), 2017. 7(8): p. 2719–2727.

29. Ewen-Campen, B., et al., Accessing the Phenotype Gap: Enabling Systematic Investigation of Paralog Functional Complexity with CRISPR. Dev Cell, 2017. 43(1): p. 6–9.

30. Hu, Y., et al., An integrative approach to ortholog prediction for disease-focused and other functional studies. BMC bioinformatics, 2011. 12(1): p. 357–357.

31. Tsherniak, A., et al., Defining a Cancer Dependency Map. Cell, 2017. 170(3): p. 564–576.e16.

32. César-Razquin, A., et al., A Call for Systematic Research on Solute Carriers. Cell, 2015. 162(3): p. 478–487.

33. Machado, J., et al., Genomic analysis of nucleoside transporters in Diptera and functional characterization of DmENT2, a Drosophila equilibrative nucleoside transporter. Physiol Genomics, 2007. 28(3): p. 337–47.

34. Xu, C., et al., An in vivo RNAi screen uncovers the role of AdoR signaling and adenosine deaminase in controlling intestinal stem cell activity. Proceedings of the National Academy of Sciences, 2020. 117(1): p. 464–471.

35. Hu, Y., et al., The Drosophila Gene Expression Tool (DGET) for expression analyses. BMC Bioinformatics, 2017. 18(1).

36. Okamoto, N., et al., A Membrane Transporter Is Required for Steroid Hormone Uptake in Drosophila. Developmental Cell, 2018. 0(0).

37. Deweirdt, P.C., et al., Accounting for small variations in the tracrRNA sequence improves sgRNA activity predictions for CRISPR screening. Nature Communications, 2022. 13(1).

38. Cherbas, L., et al., The transcriptional diversity of 25 Drosophila cell lines. Genome research, 2011. 21(2): p. 301–14.

39. Schneider, I., Cell lines derived from late embryonic stages of Drosophila melanogaster. J Embryol Exp Morphol, 1972. 27(2): p. 353–65.

40. Oberhofer, G., T. Ivy, and B.A. Hay, Cleave and Rescue, a novel selfish genetic element and general strategy for gene drive. Proceedings of the National Academy of Sciences, 2019. 116(13): p. 6250–6259.

41. Champer, J., et al., A toxin-antidote CRISPR gene drive system for regional population modification. Nature Communications, 2020. 11(1).

42. Marino, N.D., et al., Discovery of widespread type I and type V CRISPR-Cas inhibitors. Science, 2018. 362(6411): p. 240–242.

43. Ren, X., et al., High-throughput PRIME-editing screens identify functional DNA variants in the human genome. Molecular Cell, 2023. 83(24): p. 4633–4645.e9.

44. Hanna, R.E., et al., Massively parallel assessment of human variants with base editor screens. Cell, 2021. 184(4): p. 1064–1080.e20.

45. Shin, S., et al., Recombinase-mediated cassette exchange-based screening of a CRISPR/Cas9 library for enhanced recombinant protein production in human embryonic kidney cells: Improving resistance to hyperosmotic stress. Metab Eng, 2022. 72: p. 247–258.

46. Kim, S.H., et al., Identification of hyperosmotic stress-responsive genes in Chinese hamster ovary cells via genome-wide virus-free CRISPR/Cas9 screening. Metabolic Engineering, 2023. 80: p. 66–77.

47. Kim, J., et al., CRISPRi with barcoded expression reporters dissects regulatory networks in human cells. 2024, Cold Spring Harbor Laboratory.

48. Kim, S.H., et al., Genome-Wide CRISPR/Cas9 Screening Unveils a Novel Target ATF7IP-SETDB1 Complex for Enhancing Difficult-to-Express Protein Production. ACS Synth Biol, 2024. 13(2): p. 634–647.

49. Hart, T., et al., Evaluation and Design of Genome-Wide CRISPR/SpCas9 Knockout Screens. G3 Genes|Genomes|Genetics, 2017. 7(8): p. 2719–2727.

